# Analysis methods for measuring fNIRS responses generated by a block-design paradigm

**DOI:** 10.1101/2020.12.22.423886

**Authors:** Robert Luke, Eric Larson, Maureen J Shader, Hamish Innes-Brown, Lindsey Van Yper, Adrian KC Lee, Paul F Sowman, David McAlpine

**Affiliations:** Macquarie University Hearing & Department of Linguistics, Australian Hearing Hub, Macquarie University, Sydney, Australia; Department of Speech & Hearing Sciences and Institute for Learning & Brain Sciences, University of Washington, Seattle, WA, USA; The Bionics Institute Melbourne; Eriksholm Research Centre, Oticon A/S; Department of Medical Bionics, The University of Melbourne; Department of Cognitive Science, Faculty of Medicine, Health and Human Sciences, Macquarie University, Sydney, Australia; Institute for Learning & Brain Sciences, University of Washington, Seattle, WA, USA

## Abstract

**Significance:** fNIRS is an increasingly popular tool in auditory research, but the range of analysis procedures employed across studies complicates interpretation of data.

**Aim:** To assess the impact of different analysis procedures on the morphology, detection, and lateralization of auditory responses in fNIRS. Specifically, whether averaging or GLM-based analyses generate different experimental conclusions, when applied to a block-protocol design. The impact of parameter selection of GLMs on detecting auditory-evoked responses was also quantified.

**Approach:** 17 listeners were exposed to three commonly employed auditory stimuli: noise, speech, and silence. A block design was employed, comprising sounds of 5-s duration, and 10–20 s silent intervals.

**Results:** Both analysis procedures generated similar response morphologies and amplitude estimates, and both also indicated responses to speech to be significantly greater than to noise and silence. Neither approach indicated a significant effect of brain hemisphere on responses to speech. Methods to correct for systemic hemodynamic responses using short channels improved detection at the individual level.

**Conclusions:** Consistent with theoretical considerations, simulations, and other experimental domains, GLM and averaging analyses generate the same group-level experimental conclusions. We release this dataset publicly for use in future development and optimization of algorithms.

## 1. Introduction

Functional near-infrared spectroscopy (fNIRS) is an increasingly popular technique (Yücel et al., 2017) employed to investigate auditory-cortical function, and provides for a unique set of qualities that make it ideal for auditory research. fNIRS devices are typically very quiet compared to functional magnetic resonance imaging (fMRI) with which it shares a similar biologically generated signal. fNIRS is unaffected by electrical or magnetic interference from hearing devices such as cochlear implants or hearing aids, all of which are either contra-indicated or generate large artifacts in fMRI as well as in electro- and magneto-encephalography (EEG and MEG, respectively). fNIRS devices are generally relatively portable and do not require participants or patients to be isolated in a shielded chamber, or to have their head-position fixed, making it well suited for use in low- or non-compliant groups, including children, the elderly, and the cognitively impaired. It therefore provides an ideal imaging modality for clinical applications.

fNIRS has been used to investigate a variety of auditory research questions and applications. A primary use has been the investigation of cortical processing of physical qualities of sound, such as intensity, and amplitude and frequency modulations, and auditory-spatial cues (Weder et al., 2020; Weder et al., 2018; Zhang et al., 2018). fNIRS has also been employed to evaluate the perceptual qualities of speech and listening effort, as well as language development in normal-hearing and hearing-impaired populations (Anderson et al., 2019; Lawrence et al., 2018; Mushtaq et al., 2019; Pollonini et al., 2014; Rovetti et al., 2019; Rowland et al., 2018; Sevy et al., 2010; Wiggins et al., 2016b; Wijayasiri et al., 2017; Zhang et al., 2020). Research questions relating to the development of auditory cortical function (Gervain et al., 2008), and cortical reorganization following impaired sensory input and subsequent rehabilitation (Anderson et al., 2017; Wiggins and Hartley, 2015) have been investigated using fNIRS, as have outcomes related to cochlear implantation (Anderson et al., 2019) and auditory pathologies such as tinnitus (Basura et al., 2018; Shoushtarian et al., 2020).

Despite this utility, however, relative to other neuroimaging modalities such as fMRI, EEG, and MEG, fNIRS has been employed only recently by hearing scientists, and considerable variability exists in the experimental designs and analysis techniques used by different researchers. This variability can make it difficult to interpret data sets, or to replicate or compare findings across studies, or between research teams. The experimental designs most commonly employed by auditory fNIRS researchers are block- and event-related designs. Experimenters must consider a range of factors in their experimental design, including the statistical power of the protocol, the duration of the experiment, and whether the design provides the flexibility to study the effect of interest (Birn et al., 2002; Friston et al., 1999; Henson, 2007; Mechelli et al., 2003). For example, an event-related design may enable an investigator to examine the response to individual words in an ongoing sentence, something not possible when employing a block design.

Here, we compare two common analysis procedures that can be applied in experiments employing a block design. Block-design experiments present a single stimulus type continuously for an extended time interval (e.g. 5 s), followed by an inter-stimulus interval (i.e., where no stimulus is presented) of sufficient duration for the hemodynamic response to return to an approximate basal level (Brockway, 2000; Rombouts et al., 1997). Although commonly employed, no consensus exists as to the most appropriate analysis procedures for this type of experimental design; new algorithms and procedures are regularly published without cross-validation or theoretical consideration.

Analysis procedures for block designs typically lie in one of two categories: averaging analysis, where the fNIRS measurement is segmented and averaged relative to the onset of the stimulus (Dawson, 1954); and general linear model (GLM) analysis, where one or more model hemodynamic responses are fitted to the entirety of the measured fNIRS signal (Cohen, 1997; and for a recent overview in the context of fNIRS see Huppert, 2016). The signal averaging approach assumes that the noise component of the measured fNIRS signal is a random process with zero mean, and unrelated to the biological signal of interest. In contrast, the GLM is capable of accounting for a more complex model of signal noise (Barker et al., 2013). Although for non-overlapping responses such as are assumed in a block design, the GLM model is reduced to a block average, suggesting that both analyses should tend to generate similar outcomes (Dale and Buckner, 1997; Santosa et al., 2019), due to the statistical properties of the fNIRS signal, GLM analysis may be a more appropriate method with which to analyze fNIRS data (Huppert, 2016). These two analysis methods have been described and evaluated for different fNIRS analysis parameters in computer simulations and behavioral motor experiments (Santosa et al., 2019; Tak and Ye, 2014), but a direct comparison has yet to be made for research investigating audition.

In general, auditory-cortical responses in fNIRS have been shown to be reliable at a group level (Wiggins et al., 2016a). Many investigations of auditory cortical function target relatively deep (relative to the skull) cortical regions such as Heschl’s gyrus, of which a typical fNIRS device might generate less than 1% specificity (Zimeo Morais et al., 2018). This low specificity makes individual-level measurements unreliable, largely due to the poor signal-to-noise ratio; the measured stimulus-evoked hemodynamic response is small compared to all other sources of bio-generated changes in the fNIRS signal. This challenge has motivated the need for a comparison of averaging and GLM analysis specifically for auditory fNIRS signals, in order to understand the influence of analysis choices when analyzing such a small signal-of-interest. Here, we investigate whether averaging and GLM style analysis applied to the same dataset generate data that support the same experimental conclusions.

Due to the statistical properties of the noise within fNIRS signals, GLM-style analysis has been suggested to be a more appropriate method with which to analyze fNIRS data (Huppert, 2016). As such, we also investigated the influence of the parameters employed in GLM analysis on the true and false detection-rates of sound-generated fNIRS responses. Of particular importance in fNIRS experiments is the separation (and possible reduction) of systemic contributions (changes in the measured fNIRS signal that are not due to the effect of neurovascular coupling) to the measured signal when estimating neural responses (Tachtsidis and Scholkmann, 2016). This has particular relevance for auditory experiments, as systemic components of fNIRS measurements have been shown to be related to the characteristics of acoustic stimuli (Shoushtarian et al., 2019).

Many approaches have been proposed to remove the influence of systemic components on the estimation of the neural response (Fabbri et al., 2004; Saager and Berger, 2005; Santosa et al., 2020; Scholkmann et al., 2014; Wyser et al., 2020). Most use specialized channels designed not to measure neural activity but the systemic response only. These channels typically have a source-detector separation of less than 1 cm, and are often referred to as ‘short’ channels. Recently, Santosa et al. (2020) concluded that including short-channel information as a regressor of no interest within a GLM analysis resulted in the most accurate estimation of the underlying neural response compared to spatial and temporal filtering, regression, and component analysis.

We therefore investigated the effect of including information from short channels on the detection of auditory fNIRS responses. Algorithms that remove systemic components have previously been evaluated and contrasted (Santosa et al., 2020; Scholkmann et al., 2014; Wyser et al., 2020), but we apply these methods specifically in the context of two commonly used auditory stimuli: speech and band-pass noise.

Speech is the primary mode for auditory communication, and is therefore widely employed in auditory experiments. Noise signals are often used to investigate basic auditory processing, as the statistical properties of the signal can be precisely controlled. These two stimuli are often contrasted to investigate language-specific processing, or combined to investigate speech processing in challenging listening environments. Both stimuli can hold an infinite number of forms; speech may contain prosodic cues or be spectrally degraded, and noise may comprise different frequency ranges, contain modulations in amplitude or frequency, or transition over time. Here, we employed two different stimuli: speech comprising three concatenated sentences in quiet, and a 400-Hz band of noise centered at 500 Hz.

We first describe the methods used to produce and present stimuli, and to generate data. We then undertake qualitative analysis examining the morphology of fNIRS responses to auditory stimuli using averaging and GLM analyses, and assess the influence of different analysis parameters on the detection of auditory fNIRS responses, and on the rate of false positives. Finally, we investigate whether the averaging and GLM approaches provide similar experimental conclusions when applied to the same dataset. Both approaches were used to investigate two common questions in auditory neuroscience. First, do two different stimulus conditions generate a different response amplitude? Second, are cortical-hemispheric difference apparent in evoked responses?

One challenge when developing an experimental protocol for fNIRS is to understand the effects of different analysis choices, and to optimize the signal-processing procedure. Further, it is important not to optimize a specific analysis pipeline using the same data from which scientific conclusions will be drawn (Kriegeskorte et al., 2009). The dataset we report here will be released publicly to assist in the development of future auditory fNIRS pipelines and algorithm development. In a similar vein, we note that that we are not endeavouring to generate scientific conclusions concerning the relative cortical processing of speech and noise. Rather, our intention is to provide an understanding of the choice of parameters on conclusions reached by statistical analysis of auditory-generated fNIRS responses generated using averaging and GLM techniques.

## 2. Methods

### 2.1 Experimental Design

Seventeen participants volunteered for this project. All participants indicated no history of hearing concerns. Participants were aged between 22 and 40 years. Data were collected under the Macquarie University Ethics Application Reference 52020640814625.

Participants were seated in a sound-attenuating booth in a comfortable chair for the duration of the experiment, which lasted approximately 25 minutes. Participants were instructed not to pay attention to the sounds and were offered the choice of watching a silent, subtitled film during the experiment; seven participants accepted this option. NIRS data were recorded using a NIRx NIRScoutX device with APD detectors. The data were saved to disk with a sample rate of 5.2 Hz. 12 source channels and 12 detector channels were employed in the fNIRS optode-cap configuration, with eight additional short detectors distributed across the head. Sources were placed at the positions AF7, F3, F7, FC5, T7, CP5, O1, POz, O2, Iz, CP6, and T8. Detectors were placed at the positions F5, C5, TP7, CP3, P5, PO3, P04, Oz, P6, CP4, TP8, and C6. Short detectors were placed at AF7, F7, T7, CP5, O1, O2, CP6, and T8 (Figure 1). These optodes were selected to target four regions of interest (ROI) using the fOLD toolbox (Zimeo Morais et al., 2018), including the left inferior frontal gyrus (IFG), the left and right superior temporal gyri (STG), and the occipital lobe. The left inferior frontal gyrus is indicated in speech and language processing, whilst the superior temporal gyri are indicated in auditory processing. The occipital lobe is indicated in visual processing and as a possible additional site for speech processing, particularly in cross-modal plasticity studies, but this region was not expected to show significant responses in the current study.

**Figure 1:**
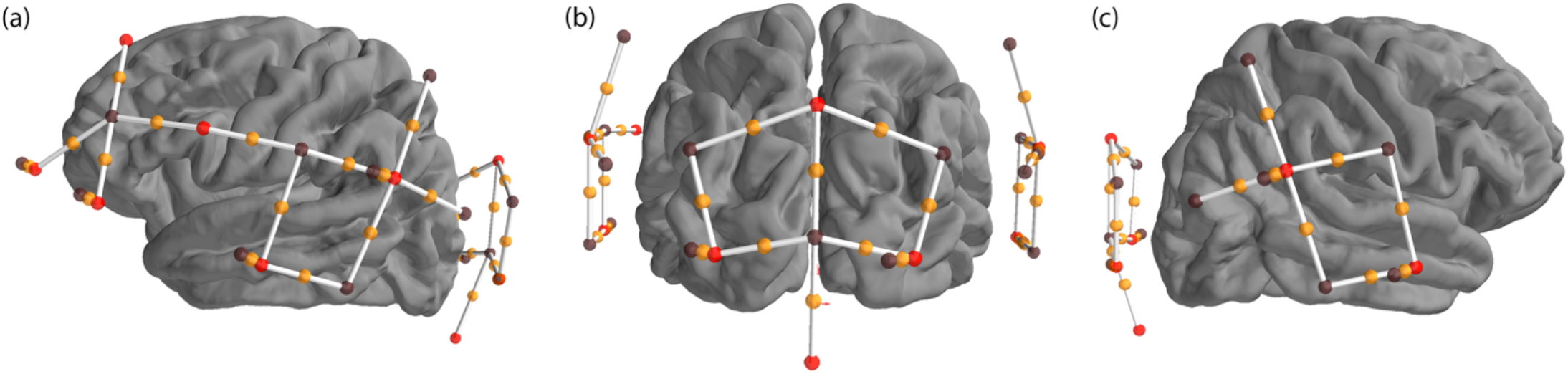
Location of sources and detectors. Four regions of interest were created to cover the left inferior frontal gyrus, the left and right superior temporal gyri, and the occipital region. Sources are shown as red dots, detectors are shown as black dots, channels are shown as white lines with an orange dot representing the midpoint. The montage is shown from the left (a), back (b) and right (c) views of the brain.

Participants listened to auditory stimuli presented diotically (i.e., the same sound to both ears) *via* Etymotic Research ER-2 insert-phones connected to an RME Fireface UCX soundcard (16 bits, 44.1 kHz sampling rate). Speech was presented at 80 dB SPL, and noise (separately) at 85 dB SPL. Stimuli were calibrated to a Casella Cel-110/2 sound source using a Norsonic sound-level meter (Norsonic SA, Norway) and an ear simulator (RA0045 G.R.A.S., Denmark).

Participants were exposed to three stimulus conditions: speech, noise, and silence. The speech stimulus consisted of three concatenated sentences from the AusTIN speech *corpus* (Dawson et al., 2013) with a total duration of 5.25 s. The noise stimulus consisted of a uniform distribution of frequency content between 300-700 Hz, and was of 5-s duration. Five seconds of silence was used as the control condition. Stimuli were presented in random order with an inter-stimulus interval selected randomly for each trial from a uniform distribution in the range 10-20 s. Twenty trials were presented for each condition, resulting in a total of 60 trials per participant.

### 2.2 Analysis

All analyses were performed using MNE (version 0.21.2) (Gramfort et al., 2013; Gramfort et al., 2014) and MNE-NIRS (version 0.0.1) (https://mne.tools/mne-nirs/), which makes extensive use of the Nilearn package (version 0.70) (Abraham et al., 2014) for GLM analysis. First, a qualitative analysis was performed to understand the morphology of the measured signal, followed by a quantitative analysis to evaluate the influence of different parameter selection on the detection of auditory responses. Finally, both the averaging and GLM analysis techniques were used to compare the response amplitude to speech vs. noise, and for relative activation in the left vs. right cortical hemispheres. All analyses were applied to the same dataset described in Section 2.1.

#### 2.2.1 The morphology of auditory responses

Hemodynamic responses vary with location on the scalp and experimental condition (Cui et al., 2011; Stoppelman et al., 2013). As such, morphology of fNIRS responses to speech and noise stimuli was investigated qualitatively using two independent procedures. The first procedure was an averaging style analysis, and the second a finite impulse response (FIR) GLM approach. Each analysis was performed on each of the three experimental conditions.

##### 2.2.1.1 Averaging analysis

The averaging analysis consisted of several steps, starting with down-sampling the data to 3 Hz, and conversion to optical density. The scalp-coupling index (Pollonini et al., 2014) was calculated for each channel between 0.7 and 1.45 Hz, and channels with an index value below 0.8 were removed. Data from each channel were then further cleaned by applying temporal-derivative distribution repair (Fishburn et al., 2019) and short-channel regression based on the nearest short channel (Saager and Berger, 2005; Scholkmann et al., 2014). Briefly, this approach to short-channel regression subtracts a scaled version of the signal obtained from the nearest short channel from the signal obtained from the long channel. The modified Beer Lambert law was then applied, with a partial pathlength factor of 0.1, converting the optical-density measurements to changes in hemoglobin concentration. Next, channels with source-detector separations outside the range 20-40 mm were excluded, followed by application of the signal-improvement algorithm based on negative correlation between oxygenated and deoxygenated hemoglobin dynamics (Cui et al., 2010). A bandpass filter was then applied between 0.01 and 0.7 Hz with a transition bandwidth of 0.005 and 0.3 Hz for the low- and high-pass edges, respectively. The data were cut into epochs from 3 s before stimulus onset to 14 s after, and a linear detrend was applied to each epoch. Epochs with a peak-to-peak difference in any channel exceeding 100 μM were then excluded. The average response *per* participant for each channel and for each condition was exported.

##### 2.2.1.2 Finite impulse response model analysis

In a second, and independent, analysis, data was entered into a GLM analysis using a deconvolution FIR model. This method makes no assumptions as to the shape of the hemodynamic response. Instead, a series of impulses following the onset of the stimulus are used as regressors to model the neural response. The morphology of the response can then be estimated by summing all the FIR components after multiplication by each component’s weight as estimated by the GLM. See Huppert (2016) and Santosa et al. (2018) for a summary of FIR and canonical approaches within the fNIRS context.

Prior to the GLM analysis, data were down-sampled to 1 Hz, and then converted to optical density. A lower sample rate was employed as the scalp-coupling index was not computed, and therefore, higher frequencies were not required. Next, channels with a source-detector separation outside the range 20-40 mm were excluded, and the modified Beer-Lambert law applied to the data, as for the averaging analysis. A GLM was then applied using a FIR model with 14 components (i.e., 14 s); this number of components was selected to ensure parity with the epoching-window approach employed in the averaging analysis. Channels were then combined into a ROI by averaging the estimates with an inverse weighting by the standard error of the GLM fit. The individual-level FIR results were then entered into a linear mixed-effects (LME) model to extract the effect of FIR delay, condition, and chromophore, whilst accounting for the random effect of subject. Santosa et al. (2018) provides for a description of these second-level statistical models.

#### 2.2.2 Canonical model analysis: Effect of parameters on response detection

Next, the effect of several analysis parameters on the detection rate for auditory responses was investigated. In contrast to the FIR approach (Section 2.2.1), this analysis used a predefined canonical model of the evoked hemodynamic response function (HRF), specifically the canonical SPM HRF, which is generated from a linear combination of two Gamma functions (Penny et al., 2011). The effect of sampling rate, correction for systemic responses, and boxcar duration on the true and false-positive detection rates was explored. For simplicity, we visualized only the data for oxyhemoglobin, and not deoxyhemoglobin, signals, as the effects of different parameters was similar for both.

Only responses from optodes placed over the superior temporal gyrus were analyzed. A false positive was defined as a response detected in the (control) condition of silence. A true positive was defined as a response detected to the speech and noise conditions. Using these definitions, a receiver operating characteristic (ROC) was defined for each analysis procedure, and the area under the curve was extracted to quantify the analysis performance. We also extracted the true positive rate (TPR) resulting from a false-positive rate (FPR) of 5%, as commonly employed in clinical studies.

Specific analysis parameters were varied in this section, but each analysis consisted of the same general procedure—a re-sampling the data, followed by conversion to optical density and hemoglobin concentration. Next, channels with source-detector separation outside the 20- to 40-mm range were excluded, as were any channels outside the superior temporal gyrus ROI. A design matrix was then constructed by creating a boxcar function based on the trigger timing, and convolving this with the SPM HRF. A GLM was performed on the data with this design matrix, including the use of a 4^th^-order auto-regressive noise model, generating channel-level data that were used to construct a ROC curve. Channel-level data were then combined into a ROI using a weighted-average procedure, in which each channel was weighted by the inverse of the standard error of the GLM. This procedure was termed the “No Correction” analysis.

To analyze the effect of different choices of processing, several modifications were made to the procedure outlined above. Different short-channel approaches were applied to correct for systemic response, including adding the mean of the short channels as a regressor to the GLM, adding the individual short channels as regressors to the GLM, as well as adding the principal components of the short channels as regressors to the GLM (adding either a subset, or all components, were investigated). These procedures were termed the “Systemic Corrected” analysis. Similarly, the effect of sample rate was investigated by down-sampling the raw signal using different rates.

#### 2.2.3 Comparison of conditions and response lateralization

Finally, a group-level analysis was performed to determine if the averaging and GLM analyses both provided the same conclusion to two research questions. First, is there a difference in response amplitude between the speech and noise stimuli? And second, is there a hemispheric difference in the response to speech stimuli? We focus on group-level analysis as this has been demonstrated to be reliable in auditory experiments (Wiggins et al., 2016a). We also investigate whether including the approach to correcting for the systemic response correction deemed most effective (see Section 2.3) modifies experimental conclusions.

##### 2.2.3.1 Averaging analysis

For the averaging analysis, the same approach was made as in Section 2.2, after which, the mean value between 5 and 7 s of the average waveform for each participant was exported for analysis by statistical testing.

##### 2.2.3.2 Canonical model analysis

For the canonical-model GLM analysis, two procedures were used; the No Correction approach and the Systemic Corrected approach, the latter of which included all principal components as regressors in the GLM to compensate for systemic responses. Both analyses used a sample rate of 0.6 Hz and a 3 s duration for the boxcar function.

##### 2.2.3.3 Statistical analysis

To summarize the dataset, results from the Systemic Corrected approach were entered into a linear mixed-effects model that accounted for condition, ROI, and chromophore with participant as a random variable. In Roger-Wilkinson notation this would be described as β ~ −1 + Condition:ROI:Chroma + (1|ID).

For each of the three analyses described above (averaging, GLM No Correction, GLM Systemic Corrected), a response estimate was exported for each participant, each condition, and each ROI. These data were then used to address two issues. First, using all channels over both left and right superior temporal gyri as a single ROI, a linear mixed-effects model was used to determine if the response to speech was different from that to noise. Participant was included as a random effect. In Roger-Wilkinson notation this is described as β ~ Condition + (1|ID). Second, a linear mixed-effects model was used to determine if the left superior temporal gyrus shows a different response amplitude to the right in the speech condition, described as β ~ ROI + (1|ID) in Roger-Wilkinson notation.

## 3. Results & Discussion

To ensure that the filter was parameterized correctly, as to remove unwanted components of the measurements and retain the frequency content of interest, the spectrum of the raw fNIRS data extracted from an example data file is plotted along with the expected hemodynamic response (Figure 2). The spectral content of the model boxcar function of the experiment convolved with a model neural response (Figure 2, red curve) indicates that the majority of the signal content is around 0.05 Hz, consistent with the average presentation rate of the experiment. The spectral content of an example measurement (Figure 2, black curve) indicates a clear signal generated by the systemic pulse rate of around 1 Hz. The filter-frequency response (Figure 2, blue) clearly retains the peak of the expected response, but excludes the low-frequency drift and high-frequency (pulse-rate) components.

**Figure 2:**
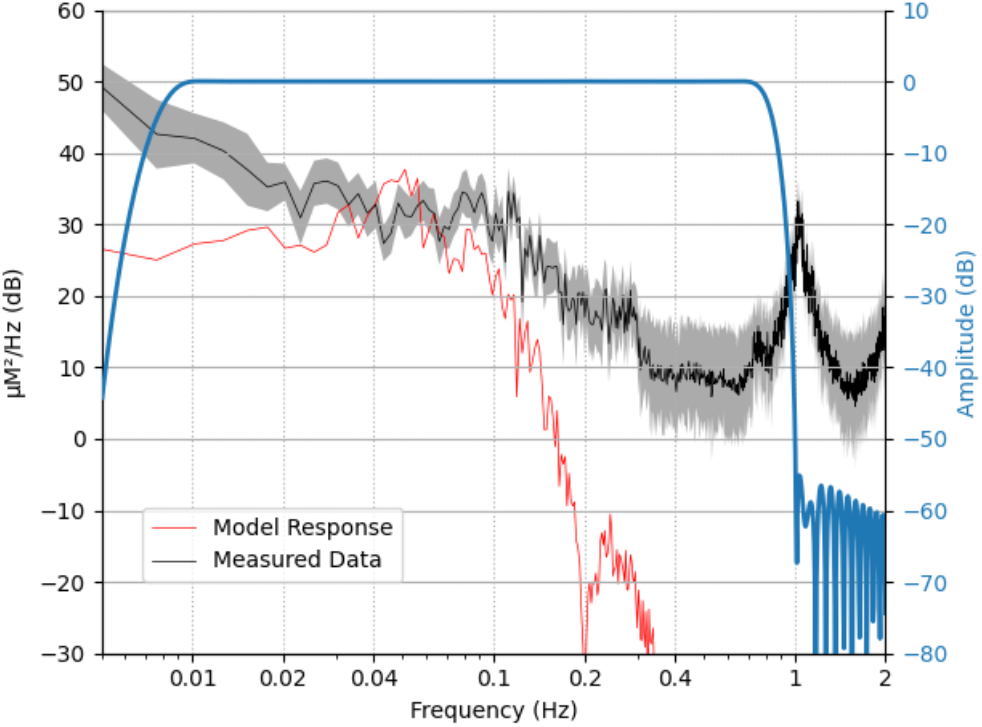
Summary of frequency information. The frequency content of the expected neural response based on trigger information and model hemodynamic response function is shown in red (arbitrary scaling). The applied filter is shown in blue. Raw data from an example file is shown in black, with the solid line indicating the mean value across all channels and the shading representing 95% confidence intervals across channels. Note that the filter retains most of the experimental frequency content while removing high-frequency heart rate content (around 1 Hz) and low frequency content in the measured data.

### 3.1 The morphology of fNIRS responses to speech and noise

Two approaches were applied to investigate the morphology of responses to auditory stimuli in each ROI. Here, we provide a qualitative description of morphology.

#### 3.1.1 Averaging analysis

To summarize the group-level averaging analysis results, a time series visualizes the average signal across participants and a bootstrapped 95% confidence band around the mean for each condition and ROI (Figure 3). Responses were observed in the STG regions for both noise and speech stimuli, but not for the silent conditions. For the silence condition, flat measurements were observed over the entire waveform in all ROIs. For both speech and noise conditions, the largest responses were measured from optodes placed over the left and right superior temporal gyri. These responses show a canonical hemodynamic response, with a peak response around 5- to 7-s after stimulus onset, consistent with the duration of the stimulus. As such, only channels over the superior temporal gyri were used subsequently to quantify response morphology.

**Figure 3:**
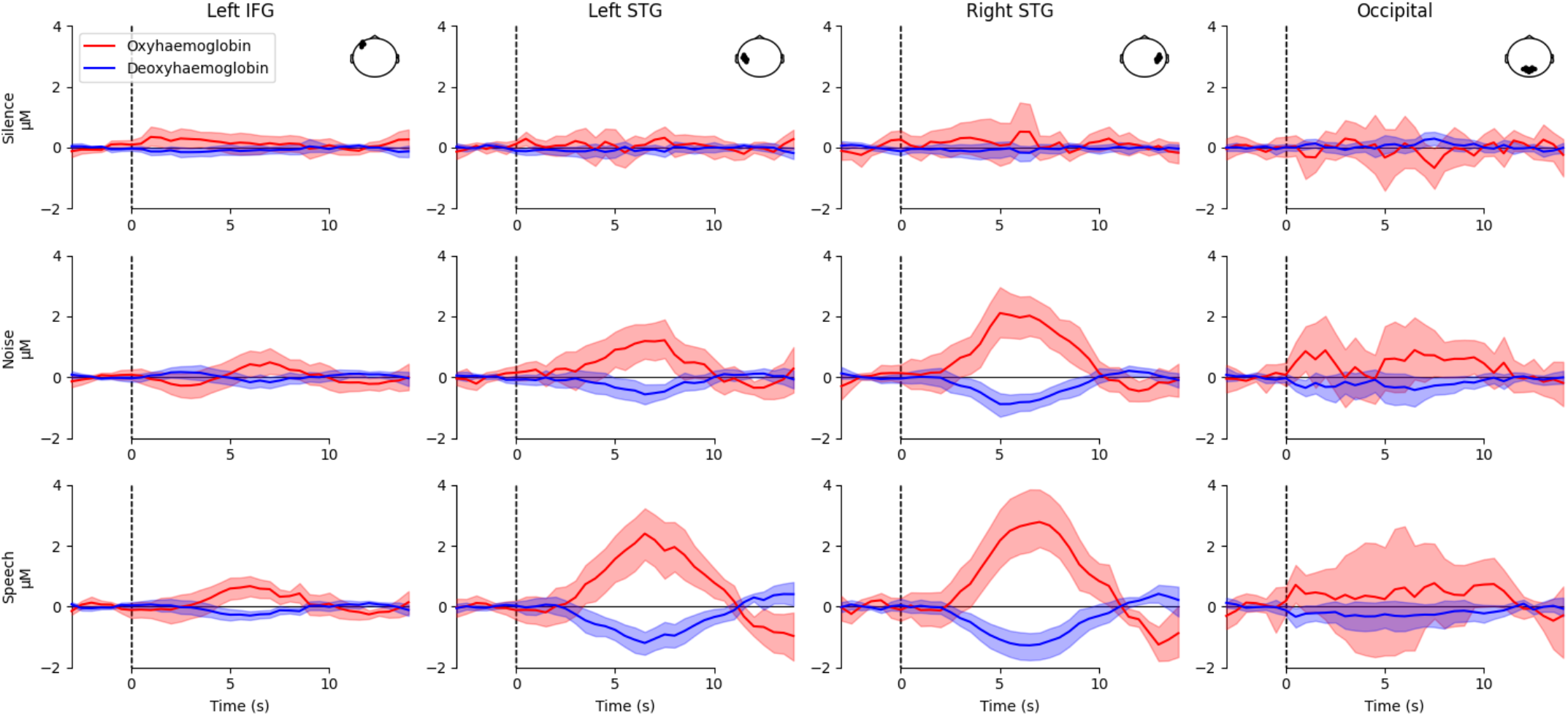
Morphology of auditory fNIRS responses using the averaging approach for all regions of interest and conditions. Each column represents a different region of interest as illustrated in the top down head view inset. Each row represents a different stimulus condition. Red represents oxyhemoglobin, blue represents deoxyhemoglobin. Shaded lines indicate 95% confidence intervals. Responses were observed over the left and right superior temporal gyrus (STG) for both speech and noise conditions, but not for silence.

#### 3.1.2 Finite impulse response model analysis

A FIR GLM analysis was also used to examine the morphology of the hemodynamic response, using only optodes situated over the superior temporal gyri. A comparison of the estimated response morphology using the averaging and the FIR (GLM) techniques (Figure 4) indicates broad agreement between the methods with regard to the timing and amplitude of hemodynamic responses, although the FIR approach generates an estimate of the response to speech greater than that suggested by averaging.

**Figure 4:**
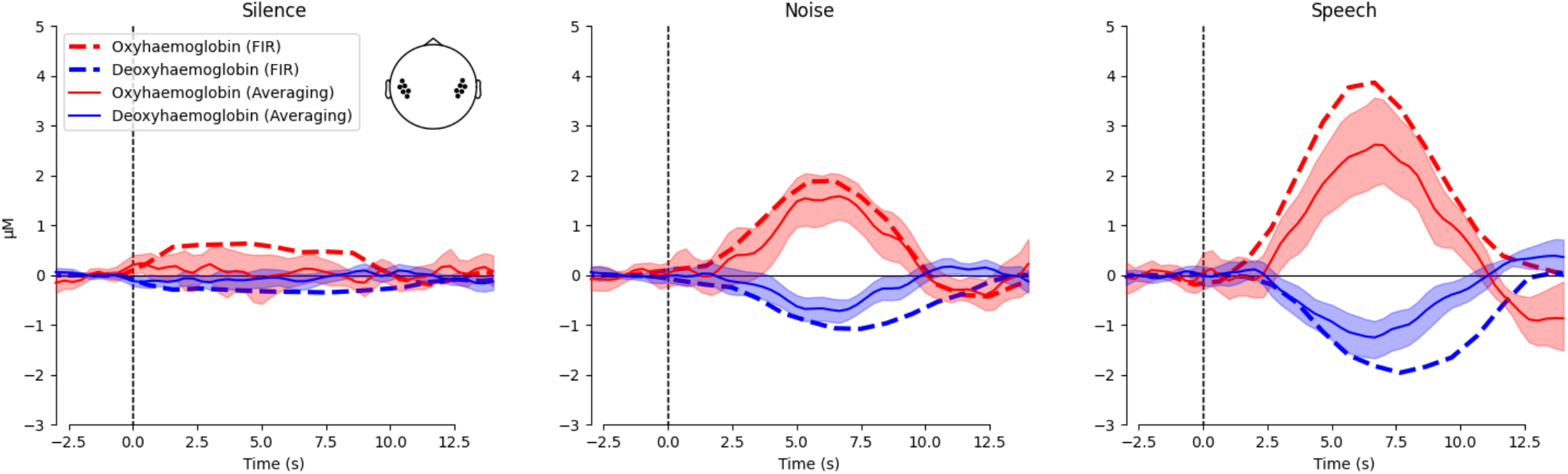
Morphology of auditory fNIRS responses over the superior temporal gyrus. Each column represents a different stimulus condition. Responses are illustrated for both oxy- and deoxyhemoglobin, red and blue respectively. The shaded areas and solid line represent the mean and 95% confidence intervals for the averaging approach. The dashed lines illustrate the estimates for the FIR GLM approach. Note that the averaging and FIR GLM fits are quite similar, except for a larger estimate for the FIR approach in the speech condition.

### 3.2 Canonical model analysis: Effect of parameters on response detection

We next examined the effect of different analysis parameters on the detection of responses in individual participants. ROC curves for both ROIs (Figure 5a) and individual channels (Figure 5b) indicates ROIs show greater sensitivity to true positives than individual channels, likely due to noisy channels being inversely weighted. Subsequently, we focus on the channel-level results (Figure 5c).

**Figure 5:**
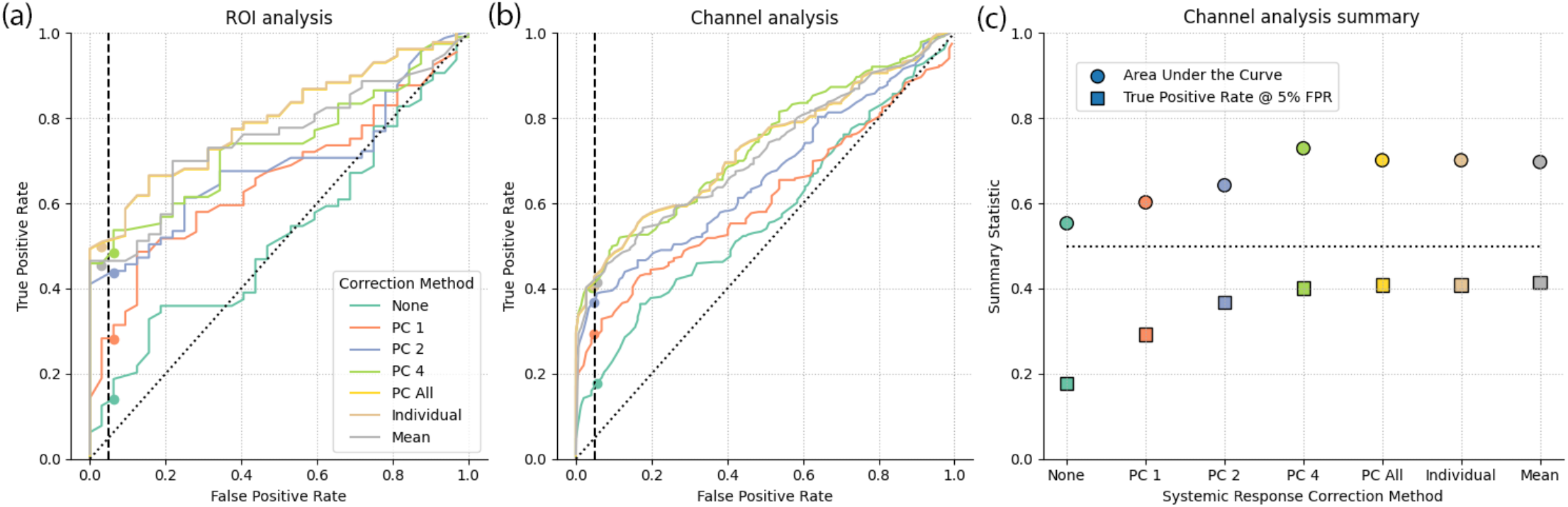
The effect of systemic response correction on auditory fNIRS response estimates. Receiver operating characteristic curves for the superior temporal gyri region of interest (a) and individual channels over the superior temporal gyri (b). Summary statistics from the individual channel ROC (c) with area under the curve (circle) and true positive rate at 5% false positive rate (square) metrics for each method. Analysis with no systemic correction is included as a reference (green), analysis with 1, 2, 4, or all principal components (PC) of the short channels as regressors in the GLM is shown (orange, blue, light green, yellow), all short channels included as individual regressors (brown) or averaged per chromophore (gray). Note that all systemic response correction approaches provide improved detection over no correction. Including all principal components, the mean of the short channels, or all individual channels provides best auditory response detection.

Two summary metrics extracted from the ROC curves are reported. First is the traditional area under the curve (AUC) measure. A larger value indicates better performance across the entire range of false positive values. Also reported is the true positive rate (TPR) occurring at the 5% false positive rate (FPR). We chose to focus on the metric at 5% FPR, as opposed to the AUC metric, because this tends to be more relevant for clinical purposes. Many of the differences in the ROC occur at a high FPR at and above 50%, however, this FPR would be considered unacceptable in a clinical setting.

#### 3.2.1 Effect of short channel regression on detection of auditory responses

We first examined the effect of different short-channel based methods of reducing systemic responses from the estimated neural responses. The effect of adding different representations of the short channels as regressors in the GLM is explored. These representations include a limited number of principal components, all principal components, the individual short channels, or the mean of the short channels per each chromophore.

Without short-channel correction, responses were detected in less than 20% of measurements for a false-positive rate of 5%. As expected, applying the short-channel method to remove systemic components resulted in a substantial improvement to the detection rate (Santosa et al., 2020; Scholkmann et al., 2014; Tak and Ye, 2014; Wyser et al., 2020). Although it is common to use just the first or second principal components as regressors (Weder et al., 2020), we observed that including all components resulted in the best performance, consistent with Santosa et al. (2020).

We also observed that including all the short channels or the mean as regressors, instead of the principal components, also results in good detection rates. Whilst we observed no effect of including all principal components or just individual channels, we selected the principal components for subsequent analysis, as this is suggested to be the most effective method to compensate for systemic components in the estimation of neural responses (Santosa et al., 2020). Neither of these approaches require a specific selection criterion, making them easy to implement, describe, and replicate.

#### 3.2.2 Effect of sample rate on the detection of auditory responses

fNIRS devices often require a trade-off between the number of channels and acquisition sample-rate, and understanding the effect of this trade-off is of practical concern for auditory experiments; performance generally decreases with lower sample rates (Figure 6). Analysis of data with a higher sample rate requires more memory and computational resources, so we selected 0.6 Hz as a sample rate that balances computational cost with accuracy.

**Figure 6:**
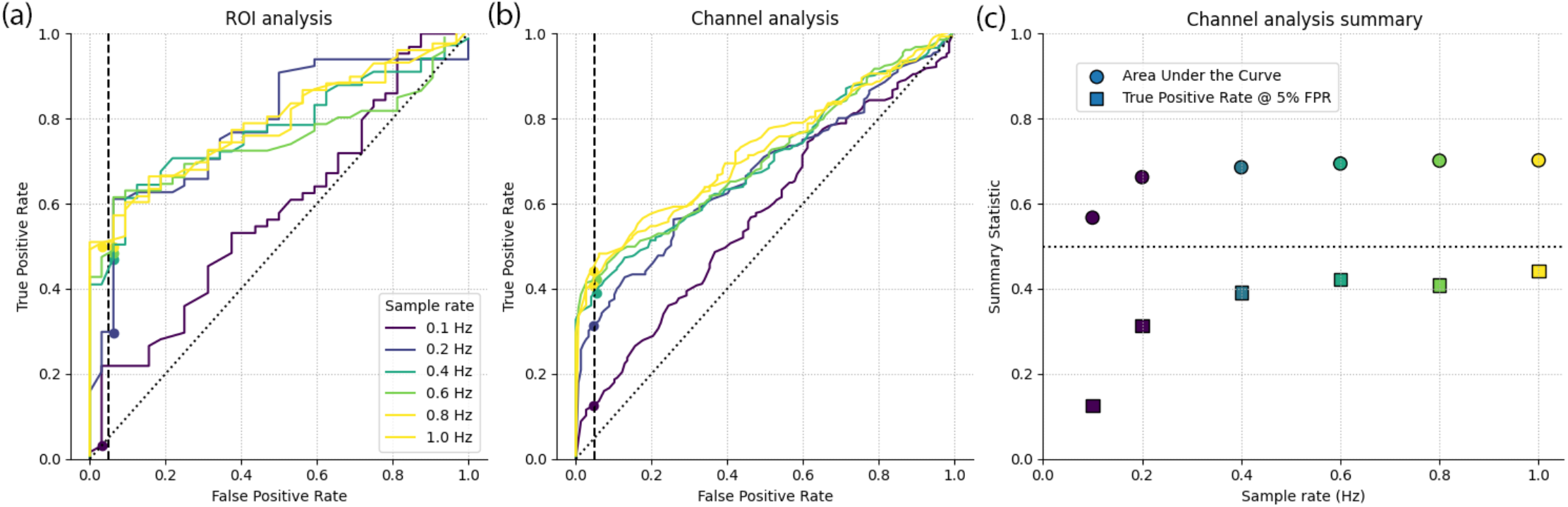
The effect of sample rate on auditory fNIRS response estimates. Receiver operating characteristic curves for the superior temporal gyri region of interest (a) and individual channels over the superior temporal gyri (b). Summary statistics from the individual channel ROC (c) with area under the curve (circle) and true positive rate at 5% false positive rate (square) metrics for data sampled at different rates. Analysis indicates improved performance with increasing sample rate, but with limited improvement above approximately 0.6 Hz.

#### 3.2.3 Effect of boxcar duration on the detection of auditory responses

The fNIRS responses to our 5-s block stimuli peaks around 6 to 7 s after stimulus onset (Figure 4). GLM analyses fit an expected neural response to the data, in which the expected neural response is generated by convolving a model HRF with a boxcar function generated from the onset times of the stimuli. The length of the boxcar function can be varied to account for the duration of the neural response, and is typically set to the duration of the stimulus. However, response morphology can change with stimuli and brain location. As such, we investigated the effect of boxcar length on response detection to auditory stimuli, and find that the 3-s boxcar function provides the greatest true positive rate, for a pre-determined 5% false-positive rate (Figure 7). Note, however, that the reduction in performance that comes from using swapping out 3-s boxcar function for one of 1-s or 5-s duration is smaller than the reduction in performance that comes about by not employing systemic correction, or when too low a sample rate is used. An alternative approach to account for differences between the model and the measured response is to include a derivative term in the design matrix (Mushtaq et al., 2020; Zhang et al., 2020). However, since we observed good correspondence between the response morphology and the expected canonical response, we did not include derivative terms in our analysis.

**Figure 7:**
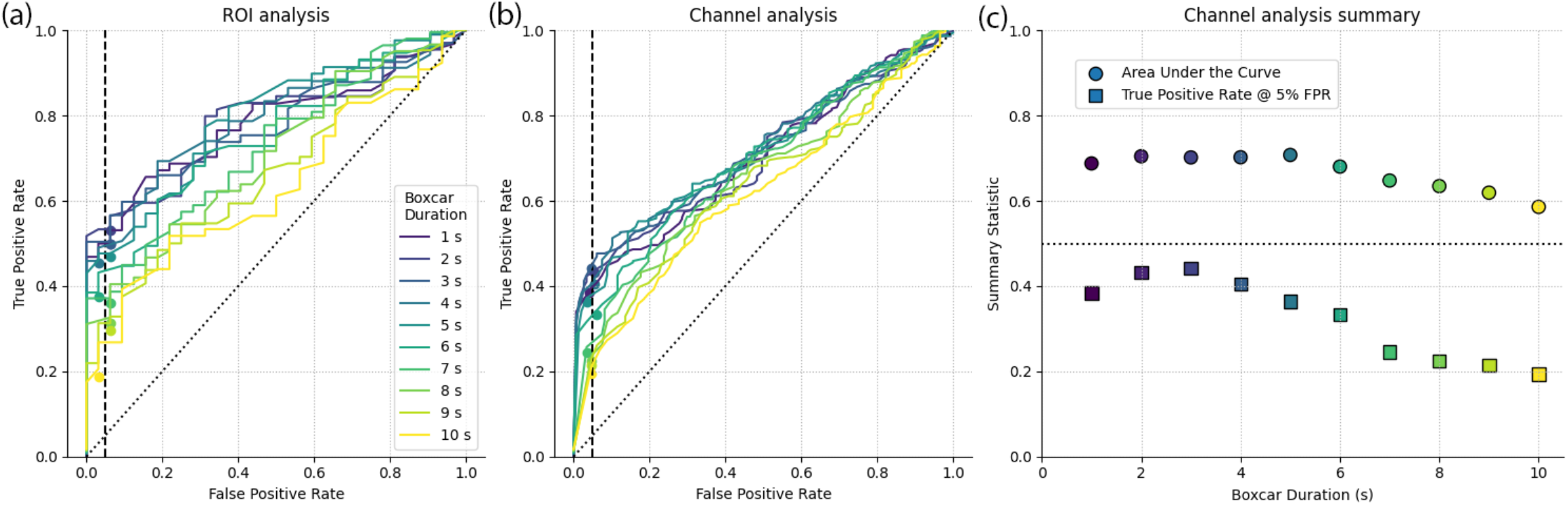
The effect of boxcar function duration on auditory fNIRS response estimates. Receiver operating characteristic curves for the superior temporal gyrus region of interest (a) and individual channels over the superior temporal gyrus (b). Summary statistics from the individual channel ROC (c) with area under the curve (circle) and true positive rate at 5% false positive rate (square) metrics for different boxcar durations. Analysis indicates optimal detection rates for a 3 s boxcar function, note that the stimulus duration was 5 s.

Additional analysis parameters beyond the scope of the current study include effects arising from selection of the specific auto-regressive model (Huppert (2016), or alternate canonical functions (Glover (1999). Based on the data thus far, we maintained a sample rate of 0.6 Hz in future analyses, and included all principal components as regressors, employing a 3-s boxcar function to model the hemodynamic response.

### 3.3 Comparison of conditions and response lateralization

Finally, we investigated whether, when applied at a group level, the averaging and GLM approaches to fNIRS analysis provide for the same experimental conclusions. Two common questions in auditory experiments were explored. First, could we detect a difference in response amplitude between two conditions, in this example: speech and noise. And second, within one condition, is a difference in response amplitudes apparent across brain hemispheres, often termed “lateralization of responses.”

We first summarized the dataset (GLM analogue of Figure 2) by modelling the response amplitude as a factor of ROI, condition, and chromophore in a LME model, with participant as a random factor (Figure 8). Consistent with the observed average waveforms (Figure 3), no significant responses were obserbed in either the left inferior frontal gyrus or occipital cortice, and the silent, control condition generated no responses in any ROI. Significant responses were observed to both speech and noise in the two ROIs of superior temporal gyrus. The lack of any detectable response to speech stimuli in left inferior frontal gyrus may be due to the passive nature of the experimental task; this cortical region has been indicated in the processing of speech, particularly in active tasks with more challenging acoustic conditions.

**Figure 8:**
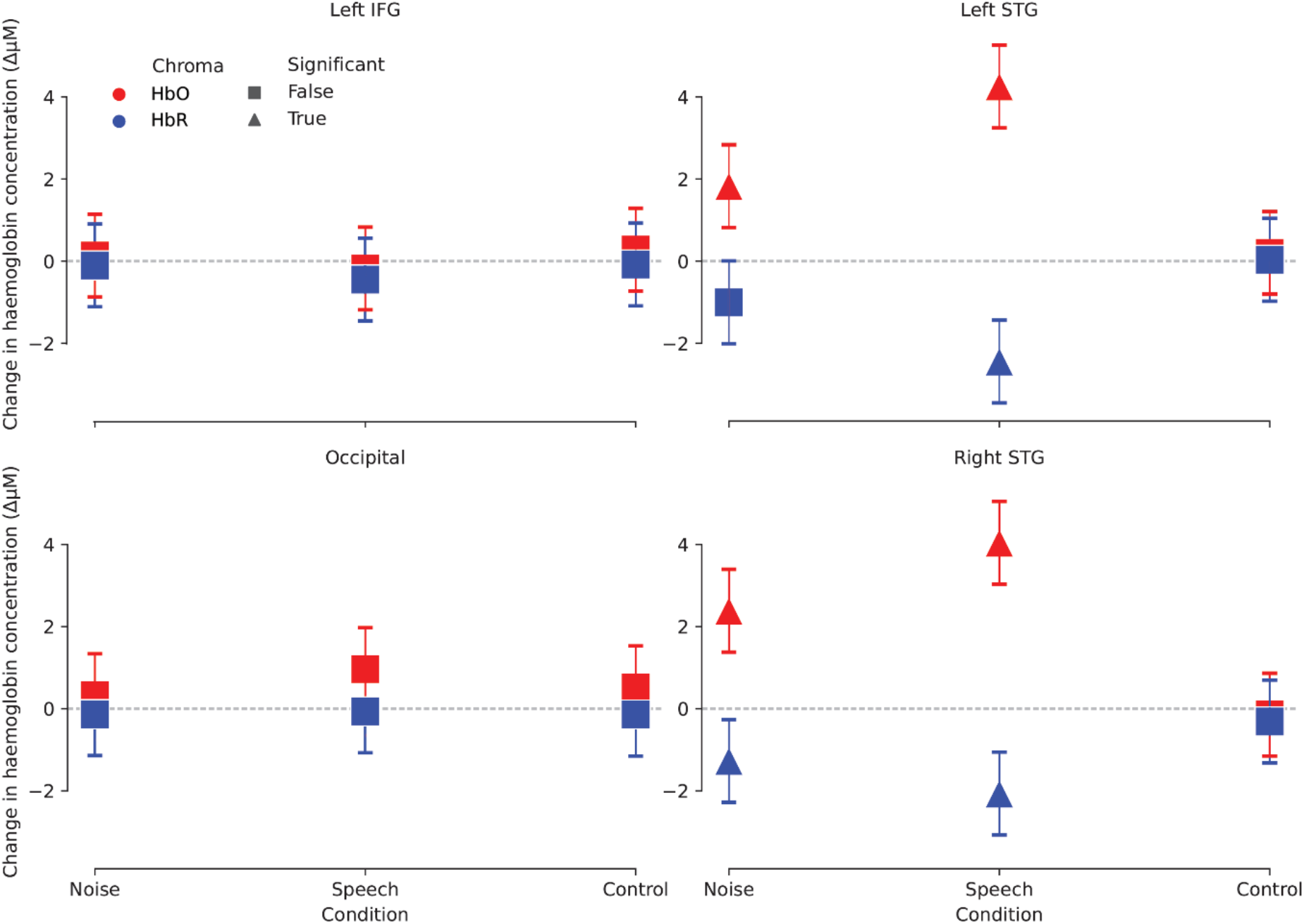
Estimates of response per condition and region of interest using the GLM analysis. Oxy- and deoxyhemoglobin responses are shown in red and blue respectively. The presence of a response (statistical difference to zero) is indicated by a triangle. Error bars represent the 95% confidence intervals of the mean.

#### 3.3.1 Does speech elicit a greater neural response than noise?

We next addressed the questions of whether responses to speech are larger than responses to noise over the superior temporal gyri ROI, and whether inter-hemispheric differences in activation are observed.

##### 3.3.1.1 Comparison of averaging and GLM result

Using the Systemic Corrected GLM with a LME model, which examined the effect of condition with participant as a random effect, we observed that the speech-evoked oxyhemoglobin response was 2.043 μM larger than that evoked by noise (*p* < .001). Using the average waveform amplitude 5 to 7 s post stimulus onset, we observed that the estimated response to speech was 1.0 μM larger than to the noise (*p* < .01). From this, we conclude that both analysis methods generate the same experimental conclusion, consistent with visual inspection of the averaging and FIR GLM analyses (Figure 3). The estimated response amplitude difference was larger for the GLM approach, possibly due to this approach better accounting for the statistical nature of the fNIRS noise (Huppert, 2016). The time window used in the averaging approach may also reduce the estimated response amplitude, whereas a peak picking approach may result in a slightly larger estimate of the response. However, automated peak-picking approaches are prone to error, particularly when the signal-to-noise ratio is low, whilst manual methods of peak-picking reduce the repeatability of an analysis.

##### 3.3.1.2 Effect of systemic component rejection

Analyzing the data using the GLM approach, with no correction for systemic responses—the No Correction analysis—indicates that the speech response was 2.306 μM larger than that to the noise stimulus (*p* = .025). Not including corrections for systemic responses generated a similar effect size to the Systemic Corrected analysis. This correspondence between methods of analysis may be due to the systemic response being relatively small, or the systemic response being similar across conditions. Our experiment was a passive listening-task, and participants were asked not to pay attention to the stimuli. Studies that have observed an event-locked systemic component to auditory stimuli required participants to generate a response, for example, by means of a button press (Shoushtarian et al., 2019). These, more-active, experimental paradigms may generate a larger systemic component, and therefore elicit greater differences between analyses corrected or uncorrected for systemic effects.

#### 3.3.2 Does speech elicit a larger response in left or right hemisphere?

##### 3.3.2.1 Comparison of averaging and GLM result

Finally, to address whether a difference in response amplitude exists between left and right cortical hemispheres to speech stimuli, results from the Systemic Corrected GLM were used in a LME model examining the effect of ROI, with participant as a random effect. The model reported that estimated amplitude of the fNIRS response in the right hemisphere was not significantly different to that in the left (β= −0.21, *p* = .73). Similarly, the same LME model reported no significant lateralization of the response amplitude when the averaging analysis was employed (β= 1.0, *p*=.13).

##### 3.3.2.2 Effect of systemic-component rejection

When assessing the No Correction GLM data at a group level, no significant effect of lateralization was observed (β= 0.18, *p*=.87), indicating that not compensating for systemic components does not generate aberrant lateralization effects. However, we cannot conclude from these data that, if a lateralization effect were present, it would be detectable without systemic correction.

## 4. Conclusion

A reference block-design auditory fNIRS dataset was created with two common acoustic stimuli. Using this dataset, it was determined that both an averaging approach and a FIR GLM analysis resulted in similar response morphology. The effect of correcting for systemic hemodynamic responses using short optical channels was evaluated on the response detection of the GLM approach, where it was determined that including the individual short channels, or the principal components of the short channels, resulted in similar practical improvements to detection. At a group level, it was observed that both the averaging and GLM approach produced the same experimental conclusions to two common research questions. Not including short-channel corrections did not change the group-level conclusions. This may be due to the fact that the task was passive in nature, and may not hold for experiments requiring active participation.

## 5. Code, Data, and Materials Availability

The fNIRS data reported in this article will be released on OSF.io and github.com in the BIDS data format to allow ease of reuse (Gorgolewski et al., 2016). All the code functions used in this analysis are available at mne.tool/mne-nirs and the associated GitHub page, along with example analysis tutorials.

## 6. Acknowledgments/Funding Sources

This study was supported by an Australian Research Council Laureate Fellowship (No. FL160100108) awarded to David McAlpine. E. Larson was supported by the National Institutes of Health under Grant R01NS104585-01A1.

## Bibliography

Abraham, A., Pedregosa, F., Eickenberg, M., Gervais, P., Mueller, A., Kossaifi, J., Gramfort, A., Thirion, B., Varoquaux, G., 2014. Machine learning for neuroimaging with scikit-learn. Front Neuroinform 8, 14.

Anderson, C.A., Wiggins, I.M., Kitterick, P.T., Hartley, D.E., 2017. Adaptive benefit of cross-modal plasticity following cochlear implantation in deaf adults. Proceedings of the National Academy of Sciences 114, 10256–10261.

Anderson, C.A., Wiggins, I.M., Kitterick, P.T., Hartley, D.E., 2019. Pre-operative brain imaging using functional near-infrared spectroscopy helps predict cochlear implant outcome in deaf adults. Journal of the Association for Research in Otolaryngology 20, 511–528.

Barker, J.W., Aarabi, A., Huppert, T.J., 2013. Autoregressive model based algorithm for correcting motion and serially correlated errors in fNIRS. Biomed Opt Express 4, 1366–1379.

Basura, G.J., Hu, X.S., Juan, J.S., Tessier, A.M., Kovelman, I., 2018. Human central auditory plasticity: A review of functional near-infrared spectroscopy (fNIRS) to measure cochlear implant performance and tinnitus perception. Laryngoscope investigative otolaryngology 3, 463–472.

Brockway, J.P., 2000. Two functional magnetic resonance imaging f(MRI) tasks that may replace the gold standard, Wada testing, for language lateralization while giving additional localization information. Brain Cogn 43, 57–59.

Cohen, M.S., 1997. Parametric analysis of fMRI data using linear systems methods. Neuroimage 6, 93–103.

Cui, X., Bray, S., Bryant, D.M., Glover, G.H., Reiss, A.L., 2011. A quantitative comparison of NIRS and fMRI across multiple cognitive tasks. Neuroimage 54, 2808–2821.

Cui, X., Bray, S., Reiss, A.L., 2010. Functional near infrared spectroscopy (NIRS) signal improvement based on negative correlation between oxygenated and deoxygenated hemoglobin dynamics. Neuroimage 49, 3039–3046.

Dale, A.M., Buckner, R.L., 1997. Selective averaging of rapidly presented individual trials using fMRI. Hum. Brain Mapping, 329–340.

Dawson, G.D., 1954. A summation technique for the detection of small evoked potentials. Electroencephalography and Clinical Neurophysiology 6, 65–84.

Dawson, P.W., Hersbach, A.A., Swanson3, B.A., 2013. An Adaptive Australian Sentence Test in Noise (AuSTIN). Ear and Hearing 34, 592–600.

Fabbri, F., Sassaroli, A., Henry, M.E., Fantini, S., 2004. Optical measurements of absorption changes in two-layered diffusive media. Phys Med Biol 49, 1183–1201.

Fishburn, F.A., Ludlum, R.S., Vaidya, C.J., Medvedev, A.V., 2019. Temporal Derivative Distribution Repair (TDDR): A motion correction method for fNIRS. Neuroimage 184, 171–179.

Gervain, J., Macagno, F., Cogoi, S., Pena, M., Mehler, J., 2008. The neonate brain detects speech structure. Proc Natl Acad Sci U S A 105, 14222–14227.

Glover, G.H., 1999. Deconvolution of impulse response in event-related BOLD fMRI. Neuroimage 9, 416–429.

Gorgolewski, K.J., Auer, T., Calhoun, V.D., Craddock, R.C., Das, S., Duff, E.P., Flandin, G., Ghosh, S.S., Glatard, T., Halchenko, Y.O., 2016. The brain imaging data structure, a format for organizing and describing outputs of neuroimaging experiments. Scientific data 3, 1–9.

Gramfort, A., Luessi, M., Larson, E., Engemann, D.A., Strohmeier, D., Brodbeck, C., Goj, R., Jas, M., Brooks, T., Parkkonen, L., Hamalainen, M., 2013. MEG and EEG data analysis with MNE-Python. Front Neurosci 7, 267.

Gramfort, A., Luessi, M., Larson, E., Engemann, D.A., Strohmeier, D., Brodbeck, C., Parkkonen, L., Hamalainen, M.S., 2014. MNE software for processing MEG and EEG data. Neuroimage 86, 446–460.

Huppert, T.J., 2016. Commentary on the statistical properties of noise and its implication on general linear models in functional near-infrared spectroscopy. Neurophotonics 3, 010401.

Kriegeskorte, N., Simmons, W.K., Bellgowan, P.S., Baker, C.I., 2009. Circular analysis in systems neuroscience: the dangers of double dipping. Nat Neurosci 12, 535–540.

Lawrence, R.J., Wiggins, I.M., Anderson, C.A., Davies-Thompson, J., Hartley, D.E.H., 2018. Cortical correlates of speech intelligibility measured using functional near-infrared spectroscopy (fNIRS). Hear Res 370, 53–64.

Mushtaq, F., Wiggins, I.M., Kitterick, P.T., Anderson, C.A., Hartley, D.E.H., 2019. Evaluating time-reversed speech and signal-correlated noise as auditory baselines for isolating speech-specific processing using fNIRS. PLoS One 14, e0219927.

Mushtaq, F., Wiggins, I.M., Kitterick, P.T., Anderson, C.A., Hartley, D.E.J.F.i.H.N., 2020. The Benefit of Cross-Modal Reorganization on Speech Perception in Pediatric Cochlear Implant Recipients Revealed Using Functional Near-Infrared Spectroscopy. 14.

Penny, W.D., Friston, K.J., Ashburner, J.T., Kiebel, S.J., Nichols, T.E., 2011. Statistical parametric mapping: the analysis of functional brain images. Elsevier.

Pollonini, L., Olds, C., Abaya, H., Bortfeld, H., Beauchamp, M.S., Oghalai, J.S., 2014. Auditory cortex activation to natural speech and simulated cochlear implant speech measured with functional near-infrared spectroscopy. Hear Res 309, 84–93.

Rombouts, S.A., Barkhof, F., Hoogenraad, F.G., Sprenger, M., Valk, J., Scheltens, P., 1997. Test-retest analysis with functional MR of the activated area in the human visual cortex. AJNR Am J Neuroradiol 18, 1317–1322.

Rovetti, J., Goy, H., Pichora-Fuller, M.K., Russo, F.A., 2019. Functional Near-Infrared Spectroscopy as a Measure of Listening Effort in Older Adults Who Use Hearing Aids. Trends Hear 23, 2331216519886722.

Rowland, S.C., Hartley, D.E.H., Wiggins, I.M., 2018. Listening in Naturalistic Scenes: What Can Functional Near-Infrared Spectroscopy and Intersubject Correlation Analysis Tell Us About the Underlying Brain Activity? Trends Hear 22, 2331216518804116.

Saager, R.B., Berger, A.J., 2005. Direct characterization and removal of interfering absorption trends in two-layer turbid media. J Opt Soc Am 22, 1874–1882.

Santosa, H., Fishburn, F., Zhai, X., Huppert, T.J., 2019. Investigation of the sensitivity-specificity of canonical- and deconvolution-based linear models in evoked functional near-infrared spectroscopy. Neurophotonics 6, 025009.

Santosa, H., Zhai, X., Fishburn, F., Huppert, T., 2018. The NIRS Brain AnalyzIR Toolbox. Algorithms 11.

Santosa, H., Zhai, X., Fishburn, F., Sparto, P.J., Huppert, T.J., 2020. Quantitative comparison of correction techniques for removing systemic physiological signal in functional near-infrared spectroscopy studies. Neurophotonics 7, 035009.

Scholkmann, F., Metz, A.J., Wolf, M., 2014. Measuring tissue hemodynamics and oxygenation by continuous-wave functional near-infrared spectroscopy--how robust are the different calculation methods against movement artifacts? Physiol Meas 35, 717–734.

Sevy, A.B., Bortfeld, H., Huppert, T.J., Beauchamp, M.S., Tonini, R.E., Oghalai, J.S., 2010. Neuroimaging with near-infrared spectroscopy demonstrates speech-evoked activity in the auditory cortex of deaf children following cochlear implantation. Hear Res 270, 39–47.

Shoushtarian, M., Alizadehsani, R., Khosravi, A., Acevedo, N., McKay, C.M., Nahavandi, S., Fallon, J.B., 2020. Objective measurement of tinnitus using functional near-infrared spectroscopy and machine learning. PLoS One 15, e0241695.

Shoushtarian, M., Weder, S., Innes-Brown, H., McKay, C.M., 2019. Assessing hearing by measuring heartbeat: The effect of sound level. PLoS One 14, e0212940.

Stoppelman, N., Harpaz, T., Ben-Shachar, M., 2013. Do not throw out the baby with the bath water: choosing an effective baseline for a functional localizer of speech processing. Brain Behav 3, 211–222.

Tachtsidis, I., Scholkmann, F., 2016. False positives and false negatives in functional near-infrared spectroscopy: issues, challenges, and the way forward. Neurophotonics 3, 031405.

Tak, S., Ye, J.C., 2014. Statistical analysis of fNIRS data: a comprehensive review. Neuroimage 85 Pt 1, 72–91.

Weder, S., Shoushtarian, M., Olivares, V., Zhou, X., Innes-Brown, H., McKay, C., 2020. Cortical fNIRS Responses Can Be Better Explained by Loudness Percept than Sound Intensity. Ear Hear 41, 1187–1195.

Weder, S., Zhou, X., Shoushtarian, M., Innes-Brown, H., McKay, C., 2018. Cortical Processing Related to Intensity of a Modulated Noise Stimulus-a Functional Near-Infrared Study. J Assoc Res Otolaryngol 19, 273–286.

Wiggins, I.M., Anderson, C.A., Kitterick, P.T., Hartley, D.E., 2016a. Speech-evoked activation in adult temporal cortex measured using functional near-infrared spectroscopy (fNIRS): Are the measurements reliable? Hear Res 339, 142–154.

Wiggins, I.M., Hartley, D.E., 2015. A synchrony-dependent influence of sounds on activity in visual cortex measured using functional near-infrared spectroscopy (fNIRS). PLoS One 10, e0122862.

Wiggins, I.M., Wijayasiri, P., Hartley, D., 2016b. Shining a light on the neural signature of effortful listening. The Journal of the Acoustical Society of America 139, 2074–2074.

Wijayasiri, P., Hartley, D.E.H., Wiggins, I.M., 2017. Brain activity underlying the recovery of meaning from degraded speech: A functional near-infrared spectroscopy (fNIRS) study. Hear Res 351, 55–67.

Wyser, D., Mattille, M., Wolf, M., Lambercy, O., Scholkmann, F., Gassert, R., 2020. Short-channel regression in functional near-infrared spectroscopy is more effective when considering heterogeneous scalp hemodynamics. Neurophotonics 7, 035011.

Yücel, M.A., Selb, J.J., Huppert, T.J., Franceschini, M.A., Boas, D.A.J.C.o.i.b.e., 2017. Functional near infrared spectroscopy: enabling routine functional brain imaging. 4, 78–86.

Zhang, M., Alamatsaz, N., Ihlefeld, A., 2020. Hemodynamic responses link individual differences in informational masking to the vicinity of superior temporal gyrus. 2020.2008.2021.261222.

Zhang, M., Mary Ying, Y.L., Ihlefeld, A., 2018. Spatial Release From Informational Masking: Evidence From Functional Near Infrared Spectroscopy. Trends Hear 22, 2331216518817464.

Zimeo Morais, G.A., Balardin, J.B., Sato, J.R., 2018. fNIRS Optodes’ Location Decider (fOLD): a toolbox for probe arrangement guided by brain regions-of-interest. Sci Rep 8, 3341.

